# Integrating memories: Congruency and reactivation aid memory integration through reinstatement of prior knowledge

**DOI:** 10.1101/716076

**Authors:** Marlieke T.R. van Kesteren, Paul Rignanese, Pierre G. Gianferrara, Lydia Krabbendam, Martijn Meeter

## Abstract

Building consistent knowledge schemas that organize information and guide future learning is of great importance in everyday life. Such knowledge building is suggested to occur through reinstatement of prior knowledge during new learning in stimulus-specific brain regions. This process is proposed to yield integration of new with old memories, supported by the medial prefrontal cortex (mPFC) and medial temporal lobe (MTL). Possibly as a consequence, congruency of new information with prior knowledge is known to enhance subsequent memory. Yet, it is unknown how reactivation and congruency interact to optimize memory integration processes that lead to knowledge schemas. To investigate this question, we here used an adapted AB-AC inference paradigm in combination with functional Magnetic Resonance Imaging (fMRI). Participants first studied an AB-association followed by an AC-association, so B (a scene) and C (an object) were indirectly linked through their common association with A (an unknown pseudoword). BC-associations were either congruent or incongruent with prior knowledge (e.g. a bathduck or a hammer in a bathroom), and participants were asked to report subjective reactivation strength for B while learning AC. Behaviorally, both the congruency and reactivation measures enhanced memory integration. In the brain, these behavioral effects related to univariate and multivariate parametric effects of congruency and reactivation on activity patterns in the MTL, mPFC, and Parahippocampal Place Area (PPA). Moreover, mPFC exhibited larger connectivity with the PPA for more congruent associations. These outcomes provide insights into the neural mechanisms underlying memory integration enhancement, which can be important for educational learning.

**Significance statement:** How does our brain build knowledge through integrating information that is learned at different periods in time? This question is important in everyday learning situations such as educational settings. Using an inference paradigm, we here set out to investigate how congruency with, and active reactivation of previously learned information affects memory integration processes in the brain. Both these factors were found to relate to activity in memory-related regions such as the medial prefrontal cortex (mPFC) and the hippocampus. Moreover, activity in the parahippocampal place area (PPA), assumed to reflect reinstatement of the previously learned associate, was found to predict subjective reactivation strength. These results show how we can moderate memory integration processes to enhance subsequent knowledge building.

## Introduction

In everyday life we continuously build up knowledge by constructing schemas that organize information and guide future learning (van Kesteren et al., 2012). Knowledge is key to success in education (Bransford et al., 2000), but it is still unclear what neural processes underlie optimal knowledge building. Reactivation of prior knowledge during learning of new, related information is suggested to aid construction of knowledge schemas (Shohamy and Wagner, 2008; Zeithamova et al., 2012b). Such neural reinstatement can yield integration of new and old memories and extend existing schemas (Zeithamova et al., 2012a; Preston and Eichenbaum, 2013; Schlichting and Preston, 2015). We recently found that active behavioral control of prior knowledge reactivation improves memory integration success (van Kesteren et al., 2018a). Therefore, to better understand knowledge building, it is important to examine how exactly reinstating prior knowledge helps to achieve memory integration.

Congruency with prior knowledge has consistently been shown to facilitate memory formation (Rojahn and Pettigrew, 1992) and is found to be supported by neural processes in the mPFC and medial MTL, specifically the hippocampus (van Kesteren et al., 2012; Gilboa and Marlatte, 2017). Previous research examining such congruency effects on memory performance has focused largely on learning episodes in which congruency was evident (e.g. van Kesteren et al., 2013a; Bein et al., 2014; Liu et al., 2018), or implicitly or explicitly learned (van Buuren et al., 2014; Brod et al., 2015; Wagner et al., 2015). However, in a recent behavioral study, we also found such a congruency effect when congruency could only be inferred through correct reactivation of prior knowledge (van Kesteren et al., 2018a). More explicitly, we showed that both congruency and subjectively reported reactivation strength independently boosted memory integration performance. These results suggest that enhancement of memory integration can be accomplished in multiple ways, governed by separate, perhaps complementary neural processes.

To investigate memory integration, an AB-AC inference paradigm is often used (Underwood, 1949; Kuhl et al., 2011; Zeithamova et al., 2012b). In this paradigm, participants first study an AB-association and a little later an AC-association. Because B and C are indirectly linked through their common association with A, participants can infer their direct relation which can lead to an integration of all three associates. Neural reinstatement of activity patterns in stimulus-specific cortex, related to the reactivated memory, is suggested to aid such integration processes (Johnson et al., 2009; Schlichting and Preston, 2016; van Kesteren et al., 2016), but it is unknown whether we can actively influence such reinstatement. Additionally, the hippocampus and mPFC are proposed to respectively encode this newly learned information and integrate it with activated prior knowledge (van Kesteren et al., 2012; Preston and Eichenbaum, 2013). Here, using fMRI, we adapted this AB-AC inference paradigm to examine reinstatement modulation. We incorporated both congruency and reactivation factors to investigate how memory integration processes in the brain can be facilitated.

In our study, participants first studied an AB-association (pseudoword-scene) followed by an AC-association (pseudoword-object), so B and C were indirectly linked through their common association with A. BC-associations were either congruent or incongruent with prior knowledge (e.g. a bathduck in a bathroom, or a hammer in a bathroom), and participants were asked to report subjective reactivation strength for B while learning AC. We expected, as we showed previously in a related behavioral experiment (van Kesteren et al., 2018a), that both congruency and reactivation would enhance memory integration. In the brain, we anticipated this to be related to neural reinstatement of scene activity (B) in the Parahippocampal Place Area (PPA) during AC-learning, showing that neural reinstatement can be actively guided. Moreover, we hypothesized that this reinstatement would be accompanied with activity in regions related to memory (MTL) and prior knowledge integration (mPFC) (van Kesteren et al., 2012; Zeithamova et al., 2012a). We investigated these hypotheses using univariate, connectivity, and multivariate analysis neuroimaging methods (Norman et al., 2006; Rissman and Wagner, 2012; van Kesteren et al., 2016).

## Materials and Methods

### Participants

We recruited 30 student participants to take part in this study. Included participants were all between 18 and 30 years old, neurologically healthy, right-handed, native Dutch speakers, and had no irremovable metal in their bodies. Recruitment was done through flyers, social media and via the participant recruitment system of the Vrije Universiteit Amsterdam (Sona Systems). In total, five participants were excluded for analyses, two of which moved their head more than the voxel size (3mm), and three participants that had low trial numbers (<10) in one or more condition that was included in the analyses. All remaining participants (n=25; 15 females) were aged 20-26 years old (mean = 21.96, SD = 1.84). They were screened for MRI safety, signed informed consent prior to participation on both study days, and were informed they could quit the experiment at any moment in time without having to give a valid reason. They were given study credits (four participants) or were paid for participation (31 participants, €38). The study was ethically approved by the ethical committees of the faculty for Behavioural and Movement Sciences of the Vrije Universiteit Amsterdam and the Faculty of Social and Behavioral Sciences of the University of Amsterdam (Spinoza centre).

### Stimuli

The stimulus set consisted of 120 object-scene associations that were either congruent or incongruent with one another (e.g. a rubber duck with a bathroom or a rubber duck with a butcher store). Congruency of associations was predefined in two counterbalanced sets such that each object and each scene belonged to either condition once (i.e. in set one the rubber duck was paired with the bathroom (congruent association) and in set two the rubber duck was paired with the butcher store (incongruent association)). Participants with an odd participant number learned set 1 and participants with an even participants number learned set 2. Additionally, we selected 60 object pictures to act as lures during the recall session. These lure objects were dissimilar from the objects used in the experiment. Stimuli were chosen to be easy to recognize, easy to define and to not have a strong emotional connotation. Scenes were four by three centimeters at 300 dots per inch (DPI) and objects (including lures) were made to fit a three by three centimeters box at 300 DPI as they had different width by length dimensions.

Next to objects and scene, we also used a set of 120 pseudowords constructed using WinWordGen (http://www.wouterduyck.be/?page_id=29, Duyck et al., 2004) which can generate pseudowords in Dutch. Pseudowords were generated under the following conditions: 4, 5 of 6 letters (evenly divided), randomly selected with 2-8 number of neighbors. Resulting words were manually checked for similarity to existing names or words. For each participant, pseudowords were randomly assigned to the object-scene associations so that behavioral effects could not be influenced by accidental familiarity with existing words.

### Procedure

When participants arrived at the scanner facility, they were welcomed by the experimenter, signed the informed consent form, and read an instruction sheet explaining the different steps of the experiment. They then performed a practice session at a computer outside the scanner to better understand the paradigm (see *Tasks* for details), and were allowed to ask questions about the procedure before going into the scanner room. They were then screened for metallic objects and were placed into the scanner. Participants laid supine on the scanner bed. A leg rest was placed under their legs and they were given an emergency alarm button in their left hand and a button box in their right hand. They were provided with ear plugs and headphones to lower the noise level of the scanner and to allow communication with the experimenter. A mirror, attached at the top of the head coil enabled participants to view the experiment screen. They were asked to move their head as little as possible during MRI image acquisition

The experiment consisted of two *encoding* runs that lasted approximately 23 and 20 minutes (8 sets and 7 sets). Between these encoding rounds participants were allowed to have a short self-paced break of a few minutes (for more details see *Tasks)*. After encoding, participants performed two *localizer* tasks that each lasted 6 minutes (for more details see *Localizer*) and at the end of the scanning session a *structural scan* that lasted four minutes was acquired. Throughout the task, participants responded using a button box consisting of four buttons of which they only used the first three. They were instructed to place the fingers of their right hand on all four buttons and use only their index, middle, and ring fingers to respond during the experiment.

At the end of the scanning session, participants left the scanner, were allowed to briefly freshen up, and were then taken to a behavioral testing room adjacent to the scanner to perform a *recall* task. They read an instruction sheet, performed a practice session, and were allowed to ask questions before the recall task was started. After the recall task, which lasted on average 45 minutes, participants went home.

Participants were invited to come back a week later to perform a second recall task and congruency rating task. They signed a second consent form, performed another practice round and completed the same recall task as the week before where items were presented in different order. Finally, they completed a congruency rating task in which they were asked to rate congruency between the objects and scenes that they learned in the scanner on a scale of 1-5 (very incongruent – very congruent; see *Tasks* for more detail), filled out an experiment-related questionnaire with questions related to strategies and difficulties, and a debriefing and payment form.

### Tasks

#### Practice

Before encoding and recall, a practice session was performed at a computer in a behavioral testing room to acquaint the participants with the experimental procedure. Participants performed the same paradigm as during the real encoding and recall tasks (see below for details), but then with only three ABC triads (one congruent and two incongruent) that were different from the ones that were included in the actual experiment. For recall, two extra lures were added. After the practice session, participants were allowed to ask questions to the experimenter or perform the practice session again until they felt they fully understood the procedure.

### Encoding

During encoding in the MRI-scanner, participants learned 120 ABC-associations (pseudoword, scene, and object). These associations were divided into 15 sets of 8 ABC triads, divided into two scanner runs of 8 sets and 7 sets respectively to allow for a brief resting period between runs. Each of these 15 sets was presented in a fixed order of tasks: AB-encoding, AB-retrieval and AC-encoding (see Figure 1). Within each of these tasks, associations were presented pseudorandomly (i.e. no more than two trials of the same category following each other), and every participant had a different predefined order.

**Figure 1:**
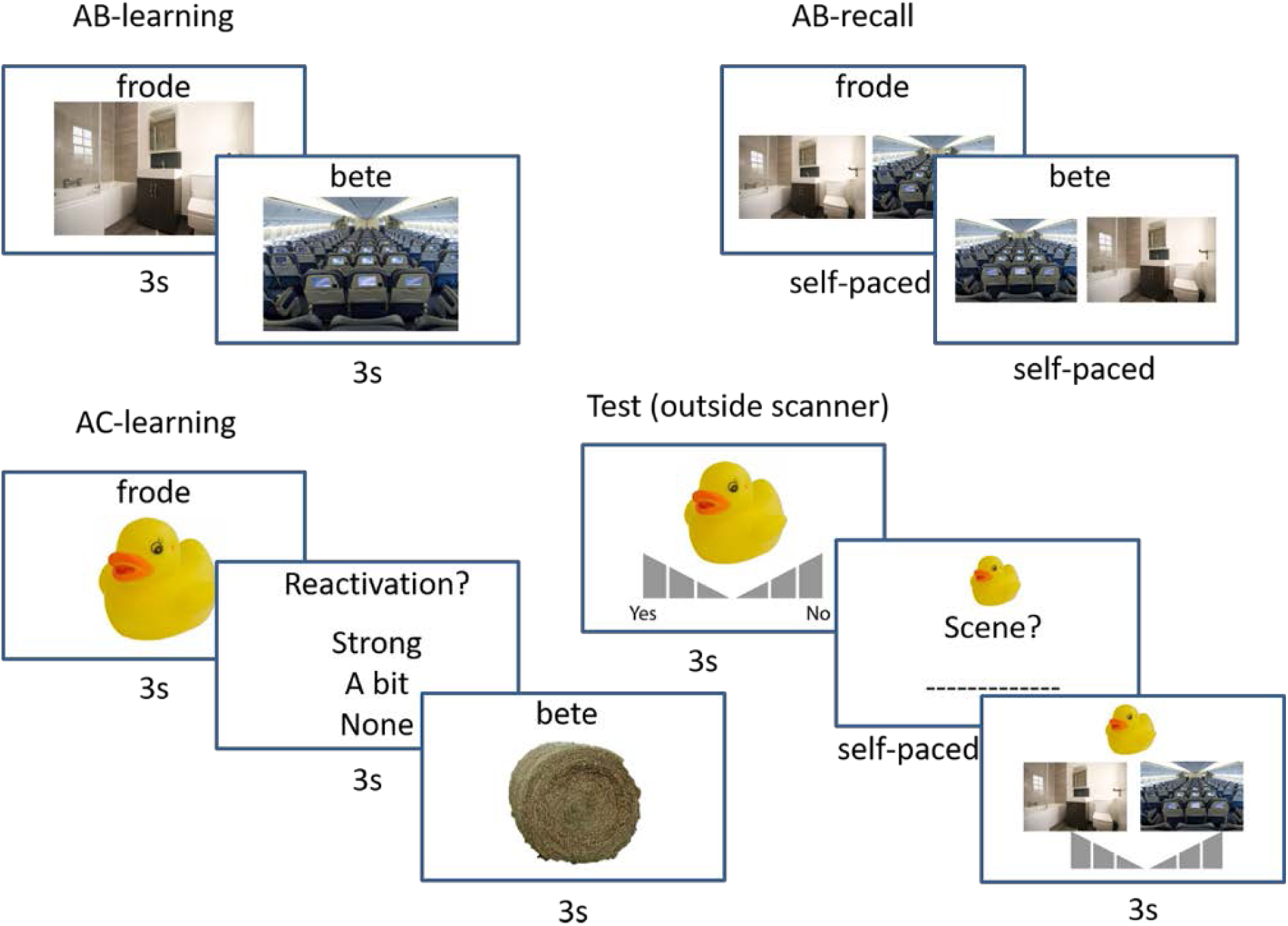
Experimental Design. Participants studied congruent and incongruent associations in an AB-AC inference paradigm (pseudoword – scene – object, e.g. frode – bathroom and frode – rubber duck or bete – airplane and bete – hay bale) in the MR-scanner. After AB-learning, a short AB-recall test was performed to further strengthen the AB-memories. After subsequent AC-learning, participants were asked how well they were able to reactivate B. After the encoding session in the scanner, participants performed a recall test in which they were asked for item recognition of the object, associative recall of the scene given the object, and associative recognition of the scene given the object. Pictures are not exactly the same as in the experiment, but very similar and rights-free.

First, participants were shown the 8 AB-associations for 3 seconds with a 5 seconds inter-trial interval (ITI) between trials, which was signaled by a “+” sign (*AB-encoding*). Every trial was initiated by the scanner pulse (TR = 2). The pseudoword was presented above the scene in the middle of the screen. Then, in order to strengthen their memory, participants completed a self-paced associative recall task (*AB-retrieval*) with a three second cut-off for all 8 AB-associations right after the end of AB-encoding. The pseudoword (A) was shown on top of the screen and participants had to select the corresponding B-item (scene) out of two possible options shown below the pseudoword. The lure scene was randomly picked from the 7 remaining scenes in the set and each scene was only presented as a lure once. Presentation side (left or right) of the correct and lure scenes was randomized. Participants were instructed to press the button corresponding to their index finger to select the left option and the button corresponding to their middle finger to select the right option. After answering, or after the cut-off of three seconds, participants immediately proceeded to the next trial, there was no ITI between these trials. AB-encoding and AB-retrieval trials combined lasted approximately 100 seconds, depending on how quick participants answered during AB-retrieval.

Third, participants completed the *AC-encoding* part of the experiment. Participants were shown AC-associations for three seconds and were instructed to try to remember the ABC-triad by reactivating the scene (B) that was associated with the same pseudoword (A). After AC-presentation, they were asked to indicate their ability to reactivate this scene using the button box: index finger: strong reactivation, middle finger: a bit reactivation, ring finger: no reactivation. They had three seconds to answer and after they answered, the ITI of 6 seconds started, indicated by a “+” sign. Since every trial was again aligned with the scanner pulse, time between presentation of sequential trials was either 12 (3+3+6) or 10 seconds (3+1+6) in cases where participants answered within one second.

### Recall

The final part of the experiment took place in a behavioral testing room adjacent to the scanner. Participants first practiced the retrieval task, with the same associations that were used during encoding practice, and were then tested on all 120 BC (scene-object) associations (see Figure 1). Participants were shown a total of 180 objects in total, 120 from the encoding task and 60 lure objects. All objects were presented in pseudorandom order such that no more than two congruent, incongruent or lure items followed each other. Order was predefined and different for each participant.

The recall task proceeded as follows: First, participants were shown an object in the middle of the screen and were given four seconds to rate their recall strength on a 6-unit Likert scale ranging from 1 - “very sure I have seen it” to 6 – “very sure I have not seen it” (object recognition). They were instructed to use the keyboard keys 1-6 to answer. After they answered, their answer option turned red and stayed on the screen for the remainder of the trial. If they did not answer in time, their answer was incorrect, or if the object was a lure, they proceeded to the next object.

If, on the other hand, the object was not a lure and they gave a correct answer, participants were asked to type a description of the associated scene (associative recall) given the object. They were instructed to be as concise and clear as possible. This test was self-paced, participants could continue to the next trial by pressing “Enter”. When they did not know the answer, they were instructed to press “Enter” without typing in an answer. Then, they were instructed to put their fingers back to the 1-6 keys (ring finger, middle finger, and index finger of both hands) and press “1”. This was added to ensure participants could answer in time on the final test.

Finally, participants were shown two scenes below the object for 3 seconds and were asked to select the scene that was indirectly associated with the object through the pseudoword (associative recognition). The other scene was randomly selected from the other scenes without replacement. They again rated their degree of confidence on a 6-unit likert scale, where 1 meant “very sure that the left scene was indirectly associated with the object”, 3 meant “poorly confident that that left scene was indirectly associated with the object”, 4 meant “poorly confident that that right scene was indirectly associated with the object” and 6 meant “very sure that the right scene was indirectly associated with the object”. The buttons 2 and 4 represented intermediate confidence for either scene. After they answered, their answer option turned red and stayed on the screen for the remainder of the trial. If they did not answer in time, they moved on to the next trial without having given an answer. The recall task lasted for 45 minutes on average. However, the amount of time spent on the task differed across participants, as some were significantly more extensive than others in answering the associative recall question. A week after the scanning session, participants performed another recall task (see *Procedure*) which was the same as the previous one, but with a different pseudorandom order.

Recall performance was scored by two independent raters that each rated the written responses to be either incorrect (0 points), partly correct (0.5 points) or correct (1 point). The partly correct answers consisted of answers that gave a vaguer description of the scene, or described part of it but went beyond general perceptual descriptions like “it was an outdoor space” or “there was a lot of blue”. Interrater similarity (Pearson correlation) was r=.96. Because associative recall for incongruent items was very low (many participants only had a few correct answers, see below), we decided to lump all memory tests together into one single *memory score* to be able to perform fMRI analyses on this measure. This memory score was calculated as a sum score for all memory tests (object recognition, associative recall, and associative recognition). Each of these tests were given 0-3 points, as both recognition tasks had a confidence rating with three options. The average score of both raters for associative recall was multiplied by 3 to give this test the same amount of weight as the recognition tests. This resulted in a memory score ranging from 0 (object recognition incorrect) to 9 (everything correct and highly confident).

### Localizer

To determine neural activity related to the processing of scenes, objects and faces, two runs of functional localizer were performed. Participants viewed blocks of 12 scenes, objects, and faces. Each stimulus was presented for 800ms with a 200ms inter-stimulus interval (ISI). At the end of each block there was a 7s ITI signaled by a “+” sign. Participants were asked to perform a one-back task and press the button corresponding to their index finger when a stimulus was shown twice. This could happen either 0, 1 or 2 times during each block and was equalized for each category, but this was not known to the participants. Responses were only checked to ensure attention to the task but were not analyzed. Importantly, none of the stimuli used in the localizer was present in the experiment. There were 30 blocks in total (10 scenes, 10 objects and 10 faces) which were divided over two runs of 15 blocks (5 scenes, 5 objects, and 5 faces) each. Order was predefined and pseudorandomized such that no block of the same category followed each other and order was the same for each participant. After the localizer, a final T1-weighted structural MRI scan lasting four minutes was acquired.

### Congruency rating

A week after the scanning session, participants were invited to come back to perform another recall task (see above) and a congruency rating task. In this task, participants were shown both the scene (on the left) and the object (on the right) and were asked to indicate how congruent they thought the association was on a scale from 1 (very incongruent) to 5 (very congruent) using keyboard buttons 1-5. This task was self-paced and associations were randomly ordered. Results were used to categorize final congruency analyses (see *Behavioral analyses* below).

### MRI acquisition procedure

Imaging scans were all acquired on 3.0T Achieva MRI system (Philips). All functional data and localizer scans were collected in 3-mm thick oblique axial slices with echo planar imaging (EPI) sequence (repetition time (TR) = 2000 ms, echo time (TE) = 28 ms; 80 × 78 × 3.7 matrix, slice thickness = 3mm; 0.3 mm gap). This resulted in approximately 1320 (on average 700 and 620) scans for encoding and 300 (2×150) scans for the localizer. At the end of the experiment, a T1-weighted structural image (256 × 256 × 172 matrix, 1 × 1 × 1.3 mm voxels) was also collected. This structural MRI image was used in data analyses for coregistration and segmentation.

### fMRI data preprocessing

Raw fMRI data from the encoding and localizer tasks were preprocessed an analyzed using statistical parametric mapping (SPM12; http://www.fil.ion.ucl.ac.uk/spm) within Matlab2014b). First, we performed motion correction using iterative rigid body realignment to minimize the residual sum of squares between the first and all further functional scans. Second, we performed rigid body coregistration to the corresponding T1 image with mutual information optimization. Subsequently, the T1 image was segmented into grey matter, white matter and Cerebrospinal Fluid (CSF), air and skull. All data were then spatially normalized through unified segmentation with bias regularisation to the Montreal Neurological Institute (MNI) 152 T1 image. For the univariate analyses, all scans were then spatially smoothed with a full-width at half maximum (FWHM) of 8mm, and for the MultiVariate Pattern Analyses (MVPA), all scans were spatially smoothed with a FWHM of 5mm and were subsequently temporally filtered with a high-pass filter (to remove scanner drift) and demeaned by computing a deviation score of the mean for each non-zero voxel in the brain and dividing by the standard deviation. MVPA-analyses were performed using Python17 as well as Nilearn and scikit-learn libraries (Abraham et al., 2014).

### Behavioral analyses

For the behavioral analyses, we calculated three measures: memory, congruency, and reactivation. See the *Recall* section above for specifics on how memory performance was calculated. The second recall test that was administered a week after scanning was not used for final analyses, because very little trials were useful. We were planning to do a memory durability analysis on these data (see Wagner et al., 2016). However, since this test was exactly the same as the first recall test, including the associative recognition phase where the right scene was shown on the screen, participants could learn associations during this test as well. Therefore, only trials that were already correctly recalled (i.e. during associative recall) were usable, yielding very little incongruent trials. We therefore decided to not focus our analyses on this test.

*Congruency* was defined as the subjective measure of congruency for each object-scene association that was indicated by the participant during the congruency rating task (see above). Congruent items were grouped as associations where participants answered “4” or “5” (congruent or very congruent) while incongruent items were grouped as associations where participants answered “1” or “2” (very incongruent or incongruent). Associations that were give a “3”, indicating that participants were indifferent about the congruency, were not taken along in this analysis. *Reactivation* was defined at the subjective measure of reactivation of the scene (B) during pseudoword-object (AC) encoding in the scanner (“Strong”, “A bit”, or “None”; see above).

### Univariate fMRI analyses

#### Localizer analyses

To determine a Region of Interest (ROI) of scene-specific activity, we created first-level models including scene, face, and object regressors along with the 6 motion regressors for each participant. We then calculated differential activity for scene > face and used these contrast maps in a second-level group analysis, calculating significant activity using a one-sample T-test. Significant activity thresholded at p < .05 Family Wise Error (FWE)-correction was then extracted to be used as a ROI in further analyses on the experimental data.

### Parametric analyses

Because of the continuous nature of our measures (memory 0-9, congruency 1-5 and reactivation 1-3), and to increase the amount of trials per condition we used parametric modulation analyses in SPM12. First-level models were constructed by adding 3 behavioral regressors, each containing trials from the different tasks during encoding: AB-encoding, AB-retrieval, and AC-encoding (see *Encoding* and Figure 1). AB-encoding and AC-encoding trials were convolved with a box-car function of 3 seconds (the full presentation time) and AB-retrieval was convolved with a box-car function of the full length of the block (maximally 24 seconds), as these trials were not separated by an ITI. For the AC-encoding trials we added our behavioral measures as four parametric modulators in the following order: memory, congruency, reactivation and reaction time, resulting in 7 behavioral regressors. Together with 6 motion regressors, and 1 continuous regressor this resulted in a model with 14 regressors per run.

The parametric modulator regressors for memory, congruency, and reactivation of the AC-encoding trials of each participant were then used in group analyses. For each of these regressors, a separate second-level one-sample T-test was performed. Because memory for incongruent items was not significantly different from chance-level, we only looked at the interactions between congruency and reactivation, which was again modelled at first-level and then again tested using a one-sample T-test at second-level.

### Connectivity analyses

To look at connectivity profiles of regions resulting from the univariate analyses, we conducted PsychoPhysiological Interaction (PPI) analyses using the Generalized PPI toolbox (gPPI), that allows for parametric PPI analyses (McLaren et al., 2012). We extracted ROIs from the univariate outcomes using SPM12 and calculated which regions were significantly influenced by an interaction between average activity in this region and the experimental variable. Based on previous PPI analyses (van Kesteren et al., 2010; van Kesteren et al., 2013a), where connectivity between the mPFC and stimulus-specific regions was found to relate to congruency, we focused specifically on connectivity profiles with the mPFC as seed. In an explorative extra analysis, we additionally considered other significantly active seeds as well. Specifics are mentioned in the results section.

### Multivariate fMRI analyses

In order to distinguish within-participant scene-related neural patterns, we trained a multivariate linear logistic regression classifier on the scene and face activity taken from the functional localizer. For feature selection, we considered activated voxels in the univariate scene > face contrast at p < .05 FWE-correction masked by a smoothed mask (5mm FWHM) of bilateral parahippocampal gyrus taken from the AAL (Automated Anatomical Labelling) template (Tzourio-Mazoyer et al., 2002). We decided to round the amount of voxels off to the participant with the lowest number of voxels in their univariate contrast, so we selected the 250 most informative voxels within this set of activated voxels for each participant (Haxby et al., 2001; Mitchell et al., 2004; Norman et al., 2006). Feature selection was performed using the “SelectKBest” function within NiBabel. To take into account the onset of the Blood Oxygenation Level Dependent (BOLD)-response, we used the localizer scans 3 TRs (6 seconds) after stimulus onset to fit our classifier (Johnson et al., 2009; Kuhl et al., 2011). Because the localizer blocks lasted for 12 seconds, we investigated two scans per series: the 4th (occurring 3 TRs after stimulus onset) and the 7th (occurring 3 TRs after the 4th scan) which resulted in 40 scans (20 scenes and 20 faces).

We trained the classifier using K-fold cross-validation by splitting the scans into 20 even chunks each chunk contained 2 scans. Importantly, scans from the same block never co-occurred in the same chunk. Using leave-one-out cross-validation, we computed performance accuracy in each fold and then computed the overall mean classifier performance accuracy. We then applied the trained classifier to our experimental data (AC-learning trials, again 6 seconds/3TRs after stimulus onset) to predict the probability of scene-related activity as opposed to face-related activity within the selected voxels. These probability values were then used to compare to behavioral measures of congruency and reactivation (see below).

### Statistics

Behavioral and neuroimaging analyses were conducted using custom codes in Python (Jupyter Notebook, https://jupyter.org) and Matlab2014b (analysis code available at https://github.com/marliekevk/Integrating-Memories). For behavioral analyses and analyses between multivariate classifier performance and behavioral measures, we used SPSS Statistics 25 (IBM) to perform one-sample T-tests to assess whether our participants performances were above chance level, and 2×3 repeated measures ANOVAs to calculate effects of congruency (congruent and incongruent) and reactivation (strong, a bit, or none) on memory performance (memory score) and classifier performance. To calculate power (Cohen’s D), we used G*Power 3.1.9.4 (Faul et al., 2007). Chance level for memory score was calculated as follows: In total, 50% of answers would be 0 (item recognition incorrect) and 50% would be > 0 (item recognition correct). For the latter cases, there are 3 (options on item recognition test) *6 (options on associative recognition test) = 18 final options, each with a 2,78% chance. This results in an average of 1.75. Given that chance level for the associative recall task is 0, this results in a global chance level of 0 + 0 + 1.75 = 1.75. For these statistical tests alpha was set at p<.05.

For the univariate fMRI analyses, results were masked to only contain grey-matter voxels. Whole-brain activity for main and interaction effects was considered significant at p < 0.05, family-wise error (FWE) correction at cluster level after an initial threshold of p < .001 uncorrected. Within predefined ROIs, either functional or structural from the AAL-template (see below for details), effects were small-volume corrected (SVC) and considered significant at p < .05 FWE-corrected at peak level after an initial threshold of p<.001 uncorrected. All coordinates mentioned are in MNI-space.

Within-participant MVPA cross-validation classifier performance was statistically compared to a null distribution which was computed using 1000 permutation tests simulating performance at chance level. To test for effects of congruency and reactivation on the classifier probabilities (indicating the degree of scene reinstatement), we performed a two-way repeated measures ANOVA (see above).

## Results

### Behavioral results

One-sample T-tests show that on average congruent (M = 3.96, SD = 1.27, t(24) = 8.64, p < .001, d = 1.74) memory scores were significantly above chance level (1.75) and incongruent (M = 2.07, SD = 1.13 t(24) = 1.40, p < .1, d = .28) memory scores were not (or at trend-level). For the combined reactivation levels, only “No reactivation” was not significantly different from chance (Strong reactivation: M = 4.36, SD = 1.52, t(24) = 8.58, p < .001, d = 1.71; A bit reactivation: M = 2.70, SD = 1.38, t(24) = 3.45, p = .001 d = .69; No reactivation: M = 2.08, SD = 1.46, t(24) = 1.15, = n.s., d = .26). When dividing results into 6 bins (congruency * reactivation), the lower reactivation bins for incongruent items were not significantly different from chance (Congruent strong reactivation: M = 5.83, SD = 1.55, t(24) = 13.16, p < .001, d = 2.63; Congruent a bit reactivation: M = 3.59, SD = 2.02, t(23) = 4.55, p < .001, d = .91; Congruent no reactivation: M = 2.42, SD = 1.90, t(24) = 1.76, p < .05, d = .35; Incongruent strong reactivation: M = 2.49, SD = 1.44, t(22) = 2.58, p < .01, d = .51; Incongruent a bit reactivation: M = 1.87, SD = 1.26, t(24) =.49, p = n.s., d = .09; Incongruent no reactivation: M = 1.75, SD = 1.51, t(24) = .01, p = n.s.., d = 0).

The repeated-measures 2×3 ANOVA contrasting effects of congruency and reactivation on memory score yielded a main effect of congruency (*F* (1,21) = 66.02, *p* < .001, η^2^ = .76), a main effect of reactivation (*F* (2,42) = 20.84, *p* < .001, η^2^ = .50), and an interaction between congruency and reactivation, (*F* (2,42) = 11.49, *p* <.001, η^2^ = .35) (Figure 2).

**Figure 2:**
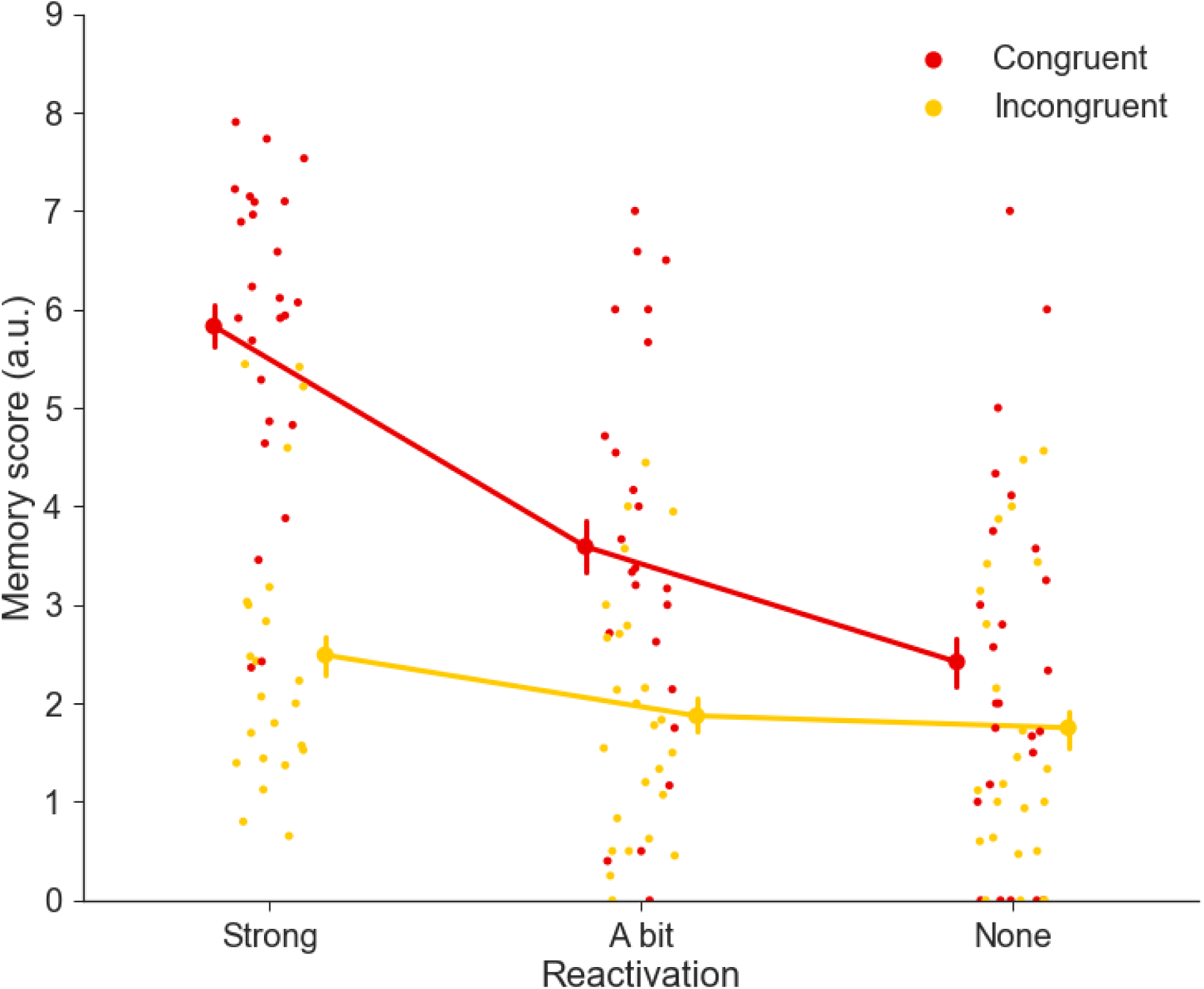
Behavioral Results. Behavioral results show main effects of congruency and reactivation on memory score, and a significant interaction between these factors. Incongruent memory performance for lower reactivation levels was not significant from chance level (1.75). Dots depict individual subject performance and error bars depict Standard Error of the Mean (SEM).

### fMRI results: univariate

All univariate results are described in Table 1 and depicted in Figure 3. For the parametric *memory* contrast, we found significant effects in dorsal visual stream extending into parahippocampal gyrus (whole-brain corrected), and bilateral anterior hippocampus (SVC-corrected with AAL bilateral hippocampus). For *congruency*, we found significant effects in bilateral parietal cortex (whole-brain corrected), left hippocampus (SVC-corrected with AAL bilateral hippocampus) and the mPFC (SVC-corrected with mPFC 10mm sphere surrounding peak [2,46,0] (van Kesteren et al., 2014)). Finally, for *reactivation*, we found significant effects in an extensive retrieval network encompassing the mPFC, and precuneus (whole-brain corrected) and the hippocampus (whole-brain corrected and SVC-corrected with AAL bilateral hippocampus to show significant overlap with hippocampus) and PPA (whole-brain corrected and SVC-corrected to show significant overlap with scene-specific activity ROI taken from the localizer). Full maps can be viewed at https://neurovault.org/collections/5673.

**Table 1:** Specifics of regions resulting from the univariate fMRI analyses. ROIs used for Small Volume Correction are described in the text. See Figure 3 for a depiction of the regions.

| Contrast | Name (AAL) | Voxels | P-value (FWE) | Peak coordinate (MNI) |
| --- | --- | --- | --- | --- |
| Memory | Middle Temporal Right | 393 | 0.025 | 50, -58, 20 |
|  | Middle Temporal Left | 1110 | < 0.001 | -44, -64, 24 |
|  | Fusiform Gyrus Right | 770 | 0.001 | 32, -40, -18 |
|  | Fusiform Gyrus Left | 848 | 0.001 | -38, -30, -18 |
|  | Hippocampus Right | 57 | 0.001 (SVC) | -22, -6, -18 |
|  | Hippocampus Left | 49 | 0.014 (SVC) | 32, -4, -20 |
| Congruency | Supramarginal Gyrus Left | 753 | < 0.001 | 60, -34, 34 |
|  | Supramarginal Gyrus Right | 453 | 0.012 | 60, -26, 24 |
|  | Hippocampus Left | 33 | 0.007 (SVC) | -28, -20, -18 |
|  | Medial Orbital Frontal Left | 19 | 0.012 (SVC) | -4, 46, -8 |
|  | Anterior Cingulate Left | 22 | 0.019 (SVC) | -6, 42, 4 |
| Reactivation | Middle Temporal Right | 5139 | < .001 | 54, -60, 18 |
|  | Angular Gyrus Right | 5148 | < .001 | -56, -60, 28 |
|  | Rectus Left | 2317 | < .001 | 0, 34, -18 |
|  | Inferior Orbital Frontal Gyrus Left | 980 | < .001 | -44, 38, -14 |
|  | Precuneus Left | 1830 | < .001 | -4, -54, 36 |
|  | Cerebellum Left | 446 | 0.005 | -32, -74, -42 |
|  | Amygdala Left | 503 | 0.003 | -24, 2, -20 |
|  | Hippocampus Left | 286 | < 0.001 (SVC) | -28, -18, -18 |
|  | Hippocampus Right | 206 | 0.003 (SVC) | 32, -20, -6 |
|  | Fusiform Right | 415 | 0.003 (SVC) | 28, -40, -8 |
|  | Fusiform Left | 63 | 0.003 (SVC) | -32, -38, -14 |
|  | Middle Occipital Left | 29 | 0.004 (SVC) | -38, -80, 22 |
|  | Middle Occipital Left | 7 | 0.008 (SVC) | -42, -78, 18 |

**Figure 3:**
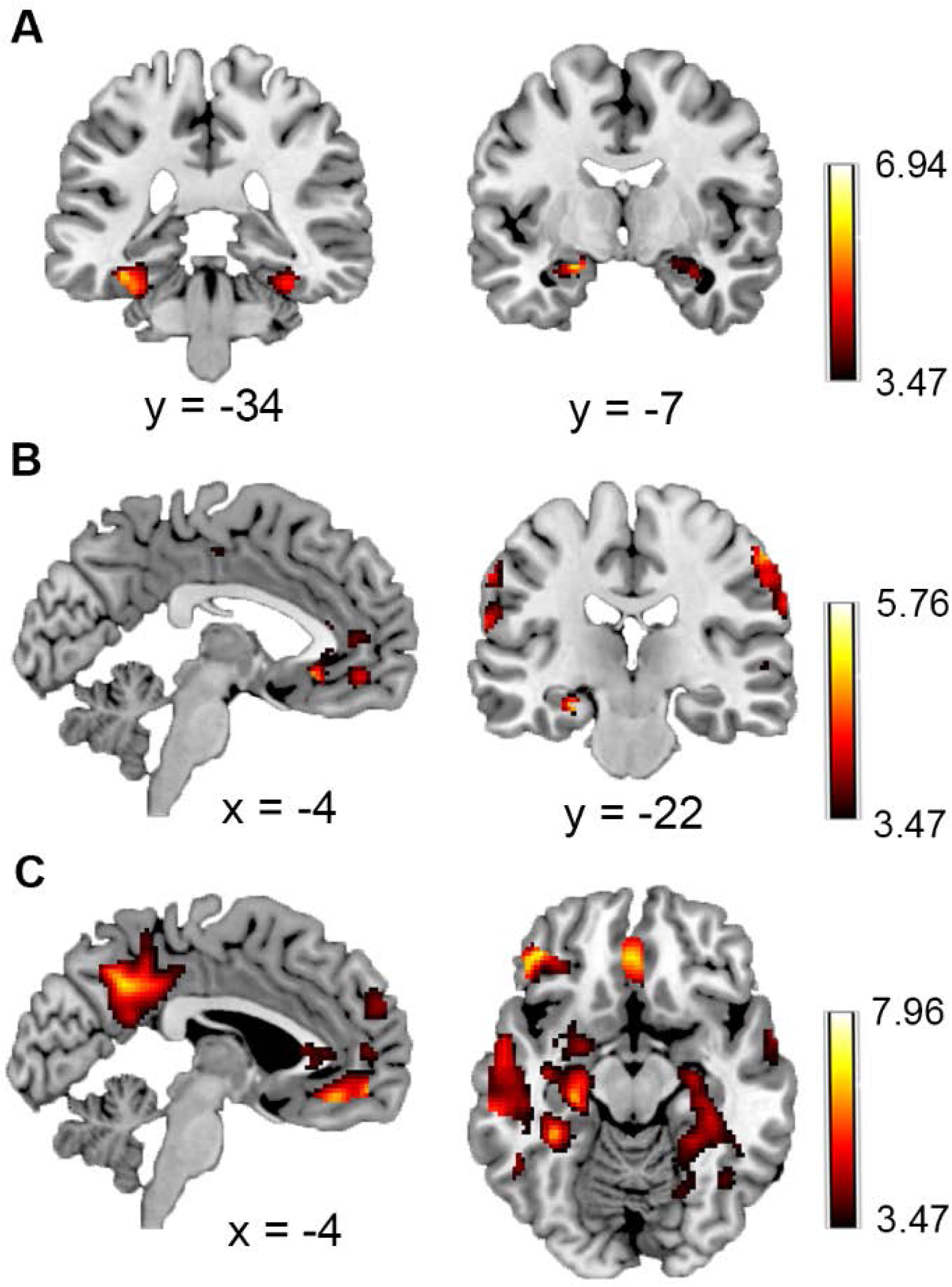
Univariate fMRI Results. Univariate fMRI results show significant effects (see Table 1) for A) *memory* in the ventral visual stream and hippocampus, B) *congruency* in the mPFC, hippocampus and parietal cortex, and C) *reactivation* in an extended retrieval network including the mPFC, hippocampus, and PPA. Maps are displayed at p < .001 uncorrected.

The opposite congruency contrast showed significant effects in Left Supplementary Motor Area (MNI [-6, 18, 48], 354 voxels, p < .05 whole-brain corrected) and Inferior Triangular Frontal Gyrus (MNI [-38, 18, 26], 257 voxels, p < .05 whole-brain corrected). The opposite reactivation contrast showed a significant effect in Right Middle Cingulate Gyrus (MNI [8,24,36], 269 voxels, p < .05 whole-brain corrected). No significant effects were found in the opposite contrast for memory and for the congruency x reactivation interaction analyses.

### fMRI results: connectivity

As mentioned in the Materials and Methods, we were mainly interested in assessing connectivity profiles between the mPFC and the rest of the brain, and possible other significantly active regions. Because the univariate contrast for reactivation yielded such a large network of activity, which would necessitate the use of many ROIs, we decided to only focus on the regions associated with the congruency contrasts. The mPFC (most significant cluster (peak [-4,46,-8]), see Table 1), was here chosen as a focus area (see Material and Methods), and we performed additional exploratory analyses on the other active regions in the congruency network (left hippocampus and bilateral parietal cortex).

PPI-results showed enhanced connectivity between the mPFC and right PPA (MNI [22, -50, -8], 24 voxels, p < .05 SVC-corrected with scene-specific activity mask taken from localizer activity) and a trend for enhanced connectivity between the mPFC and left PPA (MNI [-30,-56,-6], 6 voxels, p = .06 SVC-corrected with scene-specific ROI taken from the localizer) (Figure 4). The opposite contrast showed no significant results. We also explored connectivity profiles using the hippocampus and parietal activity clusters as seeds, but these ROIs did not show any significant connectivity as a function of congruency.

**Figure 4:**
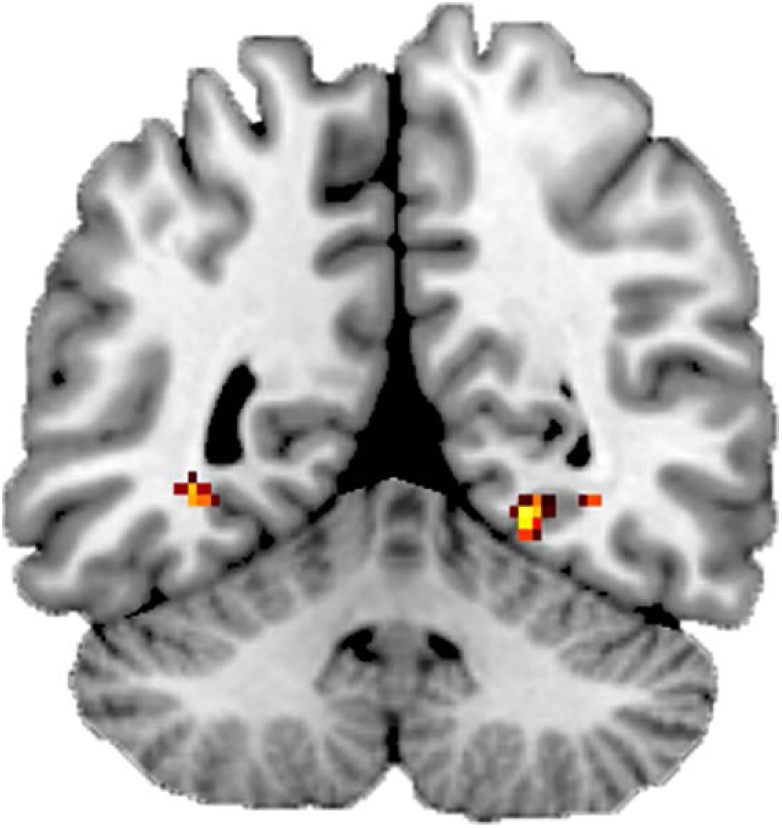
Connectivity fMRI Results. Connectivity (PPI) between mPFC and PPA was related to congruency strength. Map is displayed at p < .001 uncorrected.

### fMRI results: multivariate

Cross-validation permutation tests yielded a chance level of 48.7% and for all participants cross-validation results were significantly above chance level (p < .01). Average cross-validation performance was 92.5% (SD: 5.52%). Applying the classifier to the experimental data and relating classifier probability values to the behavioral measures for congruency and reactivation yielded a significant main effect of reactivation (F (2,42) = 6.25, p < .01, η^2^ = .10), no effect of congruency (F (1,21) = 1.33, p = n.s., η^2^ = .20), and no interaction between congruency and reactivation (F (2,42) = 1.10, p = n.s., η^2^ = .05) (Figure 5).

**Figure 5:**
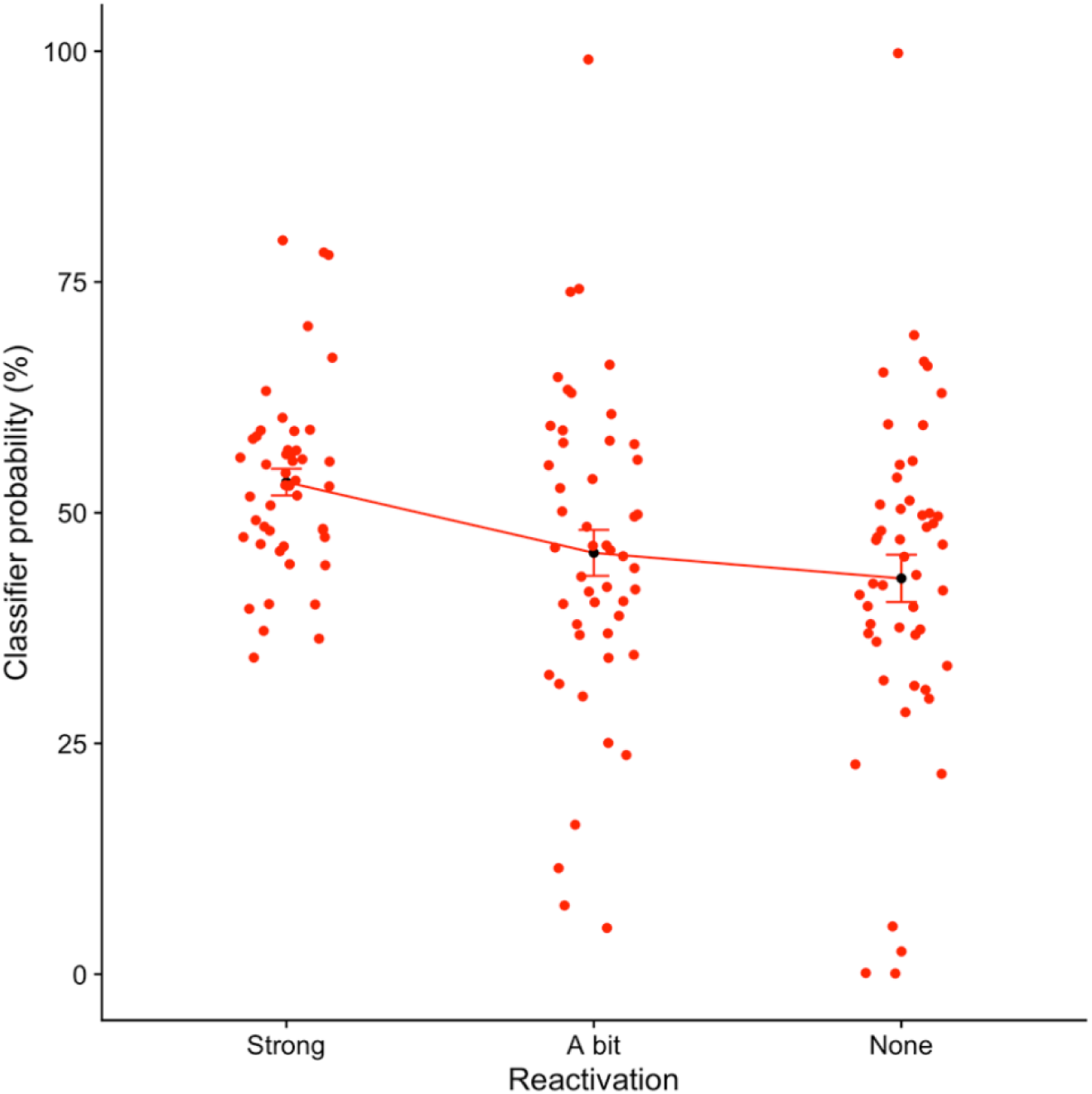
Multivariate fMRI Results. A logistic regression MVPA classifier trained on scene-specific activity showed a significant relationship to reactivation strength (p < .01). Dots depict individual subject probabilities and error bars depict Standard Error of the Mean (SEM).

## Discussion

Here, we examined neural processes underlying effects of congruency with prior knowledge, and active prior knowledge reactivation on memory integration. Using an adapted AB-AC inference paradigm, we replicated previous behavioral enhancement effects of these factors on memory performance (van Kesteren et al., 2018a). In the brain, using fMRI-univariate and multivariate analyses techniques, we show that both these factors are associated with activity profiles in the hippocampus, mPFC, and stimulus-specific activity in the PPA. Scene-related activity in the PPA, which is likely to be related to reinstatement of the reactivated scene memory during new learning, predicted reactivation strength both in univariate and multivariate analyses. Moreover, congruency was associated with increased connectivity strength between the PPA and the mPFC.

Behaviorally, we replicated our previous finding that both congruency and reactivation boost memory performance (van Kesteren et al., 2018a), this time with more abstract, non-educational associations. Additionally, we here find a significant interaction between congruency and reactivation factors as well. However, this might be related to the fact that incongruent memories were more poorly remembered overall which could be due to several factors, such as the learning environment and stimulus difficulty (pseudowords). Moreover, since we did not find such interactions in the brain, we think this interaction should be interpreted with caution.

In the brain, as expected, we show that congruency between scenes and objects related to activity in the mPFC and the hippocampus. This is consistent with previous research (Gilboa and Marlatte, 2017), but now in a task where congruency was not ubiquitous but could only be discovered through active reactivation of the scene during learning of the object. According to prevailing theories, the mPFC is suggested to aid integration of congruent associations (van Kesteren et al., 2012; Schlichting and Preston, 2015), which fits this pattern of results. Yet, not all studies show both the mPFC and the hippocampus to relate to congruency (van Kesteren et al., 2013a; van Kesteren et al., 2014), which could be related to e.g. spatial features of the learned information (van Buuren et al., 2014; Sommer, 2017; van Kesteren et al., 2018b). In this paradigm, this hippocampal activity could also reflect active retrieval of the congruency value. We additionally found enhanced connectivity between mPFC and PPA to associate to larger congruency values. This suggests that not the reinstated scene-related activity per se, but the coupling between the mPFC and the reinstated memory is more pronounced when a scene and object are congruent (van Kesteren et al., 2013b).

We found a large network of brain regions, including the mPFC, hippocampus, and PPA to relate to active reactivation during learning of new information. These regions are part of a network that is consistently found to be related to retrieval of previously learned information (Rugg and Vilberg, 2013). Moreover, decoding performance of a multivariate classifier trained on scene-specific activity within the PPA predicted reactivation strength, indicating that subjectively reported reactivation strength relates to neural reinstatement of the associated scene. Reactivation of previously learned information thus aids memory reinstatement and integration, presumably through an interaction between retrieval, encoding, and integration processes (Schlichting et al., 2015; Richter et al., 2016).

As described above, we found overlapping brain regions to relate to both congruency and reactivation factors. This suggests that differential enhancement of memory integration processes relies, at least in part, on the same network of regions. The PPA, which we anticipated to be active for both congruency and reactivation, was specifically related to reactivation strength. Yet, for congruency, this region also exhibited enhanced connectivity with the mPFC. Although the setup of our analyses (congruency was added as a parametric modulator before reactivation) should in part control for intersecting activity by accounting for factor-related variance in brain activity, we still found a lot of residual activity that correlated with reactivation strength. Moreover, interaction analyses did not yield significant results. Therefore, this pattern of results reveals an important role for both congruency and reactivation to facilitate memory integration, but it is still uncertain what the relative and complementary roles of these factors are.

Outside of our predefined ROIs, we also found significant activity in other regions. For reactivation, many regions in the dorsal and ventral visual streams were activated, which is likely related to retrieving previously learned information. Spectifically, regions in the parietal cortex were found to be related to our task as well (Wagner et al., 2005; Cabeza et al., 2012). For congruency, bilateral supramarginal gyrus was activated, and for reactivation the precuneus was activated. These regions are often found to be activated during memory retrieval (Rugg and Vilberg, 2013; Sestieri et al., 2017). Moreover, the supramarginal gyrus is adjacent to the temporo-parietal junction (TPJ) that has been indicated to be involved in schema-related processing (Gilboa and Marlatte, 2017). Previous research on schema effects on memory also showed related activity in the Angular Gyrus (Wagner et al., 2015; van der Linden et al., 2017), but we did not find such activity patterns in this study.

Every experimental approach has shortcomings and yields lessons for future research. In our experiment, we were faced with low memory performance for the incongruent condition, while in a previous educational study using the same paradigm (van Kesteren et al., 2018a), we did find significant memory for such associations. In our behavioral experiment, the main results focused on performance on the associative recall test. Here, performance on that particular test was very low for the incongruent items (i.e. participants were very poor in recalling the incongruent scenes that related to the presented object) yielding very little trials for analysis and making it hard to directly compare results. This could be related to the larger amount of associations, the setting in the MRI-scanner, and the fact that in our paradigm an abstract pseudoword was used. Therefore, for the current study, we decided to combine all memory tests into a common memory score to be able to run fMRI-analyses, and to not consider any interactions with memory performance.

Just as for memory, we also had continuous, subjective measures for congruency and reactivation, the latter two with an odd number of variables. We therefore decided to run parametric tests on our univariate fMRI data. Such tests show a correlative rather than a contrast pattern and are generally less powerful than contrasting factors. Nevertheless, we found strong results for all 3 factors, strengthening our trust in this approach. The order of parametric modulators in the first-level model can slightly affect results because a previous modulator will take away variance for next modulator. Because we were interested in effects of congruency and (most prominently) reactivation independent of memory performance, we decided to first add memory, then congruency and then reactivation to the model. However, when checking whether shifting these regressors had any effects, differences were very small so we cannot conclude that all variance related to memory and congruency is factored out with this analysis.

Finally, we did not find any interaction effects between congruency and reactivation in the brain, even though behaviorally we did see such effects. However, as described above, since incongruent items were poorly remembered, specifically for the lower reactivation values, neural interaction effects would have been difficult to interpret. Nevertheless, we are aware that the main effects reported in this paper do not control for effects of the other factors and hence we cannot conclude that we are looking at independent or differential processes.

In line with these limitations, we suggest future research to consider a paradigm in which associations are differentially reactivated, either consciously or unconsciously. Our results suggest that this can boost the neural processes underlying reinstatement and memory integration. Finding the best way to reinstate the neural patterns related to a previously learned association is expected to yield valuable practices to improve memory integration and subsequent knowledge building (van Kesteren et al., 2018a). Additionally, such research should consider developmental differences for different age groups such as children and adolescents (Brod et al., 2013), effects on long-term knowledge building and possible roles of memory consolidation (Dudai, 2012; Tompary and Davachi, 2017) and stimulus repetition (van Buuren et al., 2014; Sommer, 2017). Finally, it is important, especially for application into educational programs, to examine whether such effects are present for different types of knowledge, such as verbal, pictorial, or abstract information.

To conclude, here we show that congruency between indirectly learned scene-object associations as well as active scene-reactivation during object learning enhances memory integration performance. Using both univariate and multivariate fMRI-analysis methods, we found these behavioral results to translate to brain regions that are associated with (schema-related) memory retrieval and integration. More specifically, both congruency and reactivation factors were governed by activity in mPFC and hippocampus. Furthermore, active reactivation of previously learned information was related to univariate and multivariate activity patterns in the PPA, likely representing reinstatement of the previously learned scene while encoding the object. These results show how congruency between, and reactivation of, previously learned information enhance memory integration performance in the brain. This can be of importance in educational situations, where such enhancement is often desired.

## Acknowledgements

We would like to thank all participants for their time, Lianne de Vries for rating the associative recall performance, the staff at the Spinoza Centre for Neuroimaging, and many colleagues for support with analyses. We also thank Laura de Herde from the University of Birmingham for help with RSA-analyses that unfortunately did not yield any significant results. The authors declare no competing financial interests. This research was funded with a Marie Curie Individual Fellowship of the EU Horizon2020 Framework Program for Research and Innovation awarded to Marlieke van Kesteren (Grant number 704506).

## References

Abraham A, Pedregosa F, Eickenberg M, Gervais P, Mueller A, Kossaifi J, Gramfort A, Thirion B, Varoquaux G (2014) Machine learning for neuroimaging with scikit-learn. Front Neuroinform 8:14.

Bein O, Reggev N, Maril A (2014) Prior knowledge influences on hippocampus and medial prefrontal cortex interactions in subsequent memory. Neuropsychologia 64:320–330.

Bransford JD, Brown AL, Cocking RR (2000) How People Learn: Brain, Mind, Experience and School. Washington D.C.: National Academy Press.

Brod G, Werkle-Bergner M, Shing YL (2013) The influence of prior knowledge on memory: a developmental cognitive neuroscience perspective. Front Behav Neurosci 7:139.

Brod G, Lindenberger U, Werkle-Bergner M, Shing YL (2015) Differences in the neural signature of remembering schema-congruent and schema-incongruent events. NeuroImage 117:358–366.

Cabeza R, Ciaramelli E, Moscovitch M (2012) Cognitive contributions of the ventral parietal cortex: an integrative theoretical account. Trends in cognitive sciences 16:338–352.

Dudai Y (2012) The restless engram: consolidations never end. Annu Rev Neurosci 35:227–247.

Faul F, Erdfelder E, Lang AG, Buchner A (2007) G*Power 3: A flexible statistical power analysis program for the social, behavioral, and biomedical sciences. Behavior Research Methods 39:175–191.

Gilboa A, Marlatte H (2017) Neurobiology of Schemas and Schema-Mediated Memory. Trends in cognitive sciences 21:618–631.

Haxby JV, Gobbini MI, Furey ML, Ishai A, Schouten JL, Pietrini P (2001) Distributed and overlapping representations of faces and objects in ventral temporal cortex. Science 293:2425–2430.

Johnson JD, McDuff SG, Rugg MD, Norman KA (2009) Recollection, familiarity, and cortical reinstatement: a multivoxel pattern analysis. Neuron 63:697–708.

Kuhl BA, Rissman J, Chun MM, Wagner AD (2011) Fidelity of neural reactivation reveals competition between memories. Proc Natl Acad Sci U S A 108:5903–5908.

Liu ZX, Grady C, Moscovitch M (2018) The effect of prior knowledge on post-encoding brain connectivity and its relation to subsequent memory. NeuroImage 167:211–223.

McLaren DG, Ries ML, Xu G, Johnson SC (2012) A generalized form of context-dependent psychophysiological interactions (gPPI): a comparison to standard approaches. NeuroImage 61:1277–1286.

Mitchell TM, Hutchinson R, Niculescu RS, Pereira F, Wang XR, Just M, Newman S (2004) Learning to decode cognitive states from brain images. Mach Learn 57:145–175.

Norman KA, Polyn SM, Detre GJ, Haxby JV (2006) Beyond mind-reading: multi-voxel pattern analysis of fMRI data. Trends in cognitive sciences 10:424–430.

Preston AR, Eichenbaum H (2013) Interplay of hippocampus and prefrontal cortex in memory. Curr Biol 23:R764–773.

Richter FR, Chanales AJH, Kuhl BA (2016) Predicting the integration of overlapping memories by decoding mnemonic processing states during learning. NeuroImage 124:323–335.

Rissman J, Wagner AD (2012) Distributed representations in memory: insights from functional brain imaging. Annu Rev Psychol 63:101–128.

Rojahn K, Pettigrew TF (1992) Memory for schema-relevant information: a meta-analytic resolution. Br J Soc Psychol 31 (Pt 2):81–109.

Rugg MD, Vilberg KL (2013) Brain networks underlying episodic memory retrieval. Curr Opin Neurobiol 23:255–260.

Schlichting ML, Preston AR (2015) Memory integration: neural mechanisms and implications for behavior. Curr Opin Behav Sci 1:1–8.

Schlichting ML, Preston AR (2016) Hippocampal-medial prefrontal circuit supports memory updating during learning and post-encoding rest. Neurobiol Learn Mem 134 Pt A:91–106.

Schlichting ML, Mumford JA, Preston AR (2015) Learning-related representational changes reveal dissociable integration and separation signatures in the hippocampus and prefrontal cortex. Nat Commun 6:8151.

Sestieri C, Shulman GL, Corbetta M (2017) The contribution of the human posterior parietal cortex to episodic memory. Nature reviews 18:183–192.

Shohamy D, Wagner AD (2008) Integrating memories in the human brain: hippocampal-midbrain encoding of overlapping events. Neuron 60:378–389.

Sommer T (2017) The Emergence of Knowledge and How it Supports the Memory for Novel Related Information. Cereb Cortex 27:1906–1921.

Tompary A, Davachi L (2017) Consolidation Promotes the Emergence of Representational Overlap in the Hippocampus and Medial Prefrontal Cortex. Neuron 96:228–241 e225.

Tzourio-Mazoyer N, Landeau B, Papathanassiou D, Crivello F, Etard O, Delcroix N, Mazoyer B, Joliot M (2002) Automated anatomical labeling of activations in SPM using a macroscopic anatomical parcellation of the MNI MRI single-subject brain. NeuroImage 15:273–289.

Underwood BJ (1949) Proactive inhibition as a function of time and degree of prior learning. J Exp Psychol 39:24–34.

van Buuren M, Kroes MC, Wagner IC, Genzel L, Morris RG, Fernandez G (2014) Initial investigation of the effects of an experimentally learned schema on spatial associative memory in humans. J Neurosci 34:16662–16670.

van der Linden M, Berkers R, Morris RGM, Fernandez G (2017) Angular Gyrus Involvement at Encoding and Retrieval Is Associated with Durable But Less Specific Memories. J Neurosci 37:9474–9485.

van Kesteren MT, Brown TI, Wagner AD (2016) Interactions between Memory and New Learning: Insights from fMRI Multivoxel Pattern Analysis. Frontiers in systems neuroscience 10:46.

van Kesteren MT, Rijpkema M, Ruiter DJ, Fernandez G (2010) Retrieval of associative information congruent with prior knowledge is related to increased medial prefrontal activity and connectivity. J Neurosci 30:15888–15894.

van Kesteren MT, Ruiter DJ, Fernandez G, Henson RN (2012) How schema and novelty augment memory formation. Trends Neurosci 35:211–219.

van Kesteren MT, Rijpkema M, Ruiter DJ, Morris RG, Fernandez G (2014) Building on prior knowledge: schema-dependent encoding processes relate to academic performance. J Cogn Neurosci 26:2250–2261.

van Kesteren MT, Beul SF, Takashima A, Henson RN, Ruiter DJ, Fernandez G (2013a) Differential roles for medial prefrontal and medial temporal cortices in schema-dependent encoding: from congruent to incongruent. Neuropsychologia 51:2352–2359.

van Kesteren MT, Beul SF, Takashima A, Henson RN, Ruiter DJ, Fernandez G (2013b) Differential roles for medial prefrontal and medial temporal cortices in schema-dependent encoding: from congruent to incongruent. Neuropsychologia 51.

van Kesteren MTR, Krabbendam L, Meeter M (2018a) Integrating educational knowledge: reactivation of prior knowledge during educational learning enhances memory integration. npj Science of Learning 3:11.

van Kesteren MTR, Brown TI, Wagner AD (2018b) Learned Spatial Schemas and Prospective Hippocampal Activity Support Navigation After One-Shot Learning. Front Hum Neurosci 12:486.

Wagner AD, Shannon BJ, Kahn I, Buckner RL (2005) Parietal lobe contributions to episodic memory retrieval. Trends in cognitive sciences 9:445–453.

Wagner IC, van Buuren M, Bovy L, Fernandez G (2016) Parallel Engagement of Regions Associated with Encoding and Later Retrieval Forms Durable Memories. J Neurosci 36:7985–7995.

Wagner IC, van Buuren M, Kroes MC, Gutteling TP, van der Linden M, Morris RG, Fernandez G (2015) Schematic memory components converge within angular gyrus during retrieval. Elife 4:e09668.

Zeithamova D, Dominick AL, Preston AR (2012a) Hippocampal and ventral medial prefrontal activation during retrieval-mediated learning supports novel inference. Neuron 75:168–179.

Zeithamova D, Schlichting ML, Preston AR (2012b) The hippocampus and inferential reasoning: building memories to navigate future decisions. Front Hum Neurosci 6:70.

